# Systematic Discovery of the Functional Impact of Somatic Genome Alterations in Individual Tumors through Tumor-specific Causal Inference

**DOI:** 10.1101/329375

**Authors:** Chunhui Cai, Gregory F. Cooper, Kevin N. Lu, Xiaojun Ma, Shuping Xu, Zhenlong Zhao, Xueer Chen, Yifan Xue, Adrian V. Lee, Nathan Clark, Vicky Chen, Songjian Lu, Lujia Chen, Liyue Yu, Harry S Hochheiser, Xia Jiang, Q. Jane Wang, Xinghua Lu

## Abstract

We report a tumor-specific causal inference (TCI) framework, which discovers causative somatic genome alterations (SGAs) through inferring causal relationships between SGAs and molecular phenotypes (e.g., transcriptomic, proteomic, or metabolomic changes) within an individual tumor. We applied the TCI algorithm to tumors from The Cancer Genome Atlas (TCGA) and identified those SGAs that causally regulate the differentially expressed genes (DEGs) within each tumor. Overall, TCI identified 634 SGAs that cause cancer-related DEGs in a significant number of tumors, including most of the previously known drivers and many novel candidate cancer drivers. The inferred causal relationships are statistically robust and biologically sensible, and multiple lines of experimental evidence support the predicted functional impact of both well-known and novel candidate drivers. By identifying major candidate drivers and revealing their functional impact in a tumor, TCI shed light on disease mechanisms of each tumor, providing useful information for advancing cancer biology and precision oncology.

**Significance statements:** Cancer is mainly caused by SGAs. Precision oncology involves identifying and targeting tumor-specific aberrations resulting from causative SGAs. TCI is a novel computational framework for discovering the causative SGAs and their impact on oncogenic processes, thus revealing tumor-specific disease mechanisms. This information can be used to guide precision oncology.

## Introduction

Cancer is mainly caused by a variety of SGAs, including, but not limited to, somatic mutations (SMs) (1,2), somatic DNA copy number alterations (SCNAs)(3,4), chromosome structure variations (5-7), and epigenetic changes (8-10). Each tumor hosts a unique combination of SGAs ranging in number from hundreds to thousands, of which only a small fraction contributes to tumorigenesis (drivers), while the rest are non-consequential (passengers). Identifying causative SGAs that underlie different oncogenic processes (11), such as metastasis or immune evasion, in an individual tumor is of fundamental importance in cancer biology and precision oncology (12-14).

Current methods for identifying cancer driver genes concentrate on finding those that have a higher than expected mutation rate in a cohort of tumor samples (15-17). Some methods focus on specific mutation sites (e.g., mutation hotspots at specific amino acids or within a 3D functional domain of a protein) that likely affect the function of those proteins encoded by the mutant genes (17-22). These mutation-centered, frequency-based models have successfully identified many major oncogenes and tumor suppressors across cancer types. However, they are limited in terms of determining the functional impact of mutations, because mutation frequency of a gene (either at the gene or at the specific amino acid level) does not directly reflect which molecular or cellular processes will be affected by the mutant gene product.

In addition to mutation-frequency-centered methods, researchers have also explored combining mutation data with transcriptomic data to detect candidate drivers. Bertrand et al. (23) and Hou et al. (24) reported methods to identify mutation events that are associated changed expression of neighboring genes in a biological network (e.g., a protein-protein interaction network), based on the assumption that such changed expression can be induced by a feedback loop in response to changed function of mutated gene, and as such, the mutation is likely functional. However, these methods cannot detect direct downstream effects (other than feedback) of a mutant gene.

Besides mutations, other SGA events affecting driver genes also contribute to cancer development, such as SCNAs (3,4,25), chromosome structure variation (5-7), and epigenetic changes (8-10). Currently, analyses of SMs, SCNAs, structure variation, and epigenetic data are usually carried out separately, with distinct statistical models for different types of data (1,2,16,26,27). Such disconnection is largely due to the lack of a unifying statistical framework that is capable of integrating diverse data. The failure of integrating diverse SGA data foregoes the advantages in terms of both statistical power and biological insights that can be gained by pooling diverse information to assess the role of a driver gene in oncogenesis.

In this study, we designed a general framework based on the principle of Bayesian causal inference (28-30), referred to as Tumor-specific Causal Inference (TCI), for estimating the causal relationships between SGAs and molecular phenotypes observed in an individual tumor (31). This causality-centered framework allows integration of different SGA events to determine whether any of them have a causal impact on molecular phenotypes. Identification of SGAs that have specific functional impact on molecular phenotypes in an individual tumor can help to differentiate candidate driver SGAs from passengers and shed light on the disease mechanism of an individual tumor. Pooling information across multiple tumors provides insights about general oncogenic processes across tumors. Finally, understanding disease mechanisms of an individual tumor can guide precision oncology.

## Results

### **TCI is a** unifying **framework for discovering functional impact of SGAs in an individual tumor.**

We designed the TCI algorithm to discover the causal relationships between SGAs and DEGs observed in an individual tumor. Specifically, given a tumor *t* hosting a set of SGAs (*SGA_SET_t_*) and a set of DEGs (*DEG_SET_t_*), TCI estimates the causal relationships between SGAs and DEGs as a bipartite Bayesian causal network (28-30) (Figure 1) and searches for the causal model *M* with a maximal posterior probability P(M|D) that is most likely given the data *D* (SGAs and DEGs), which consists of data about tumor *t* and about the tumors in a set of training cases.

**Figure 1.**
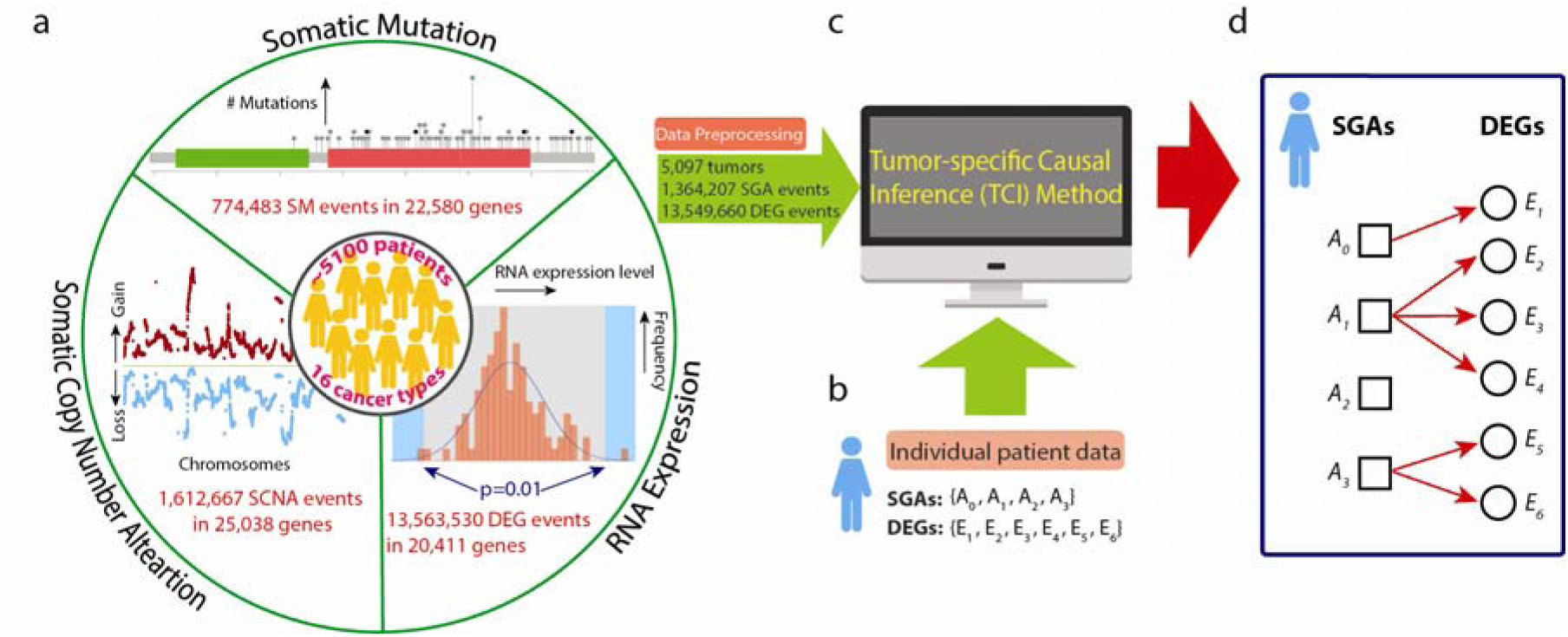
Workflow of TCI analysis, **a.** A compendium of cancer omics data is used as the training dataset. Three types of data from the 5,097 pan-cancer tumors were used in this study, including SM data (774,483 mutation events in 22,580 genes), SCNA data (1,612,667 copy number alteration events in 25,038 genes), and gene expression data (13,563,530 DEG events in 20,411 genes). SM and SCNA data were integrated as SGA data. Expression of each gene in each tumor was compared to a distribution of the same gene in the “normal control” samples, and, if a gene’s expression value was outside the significance boundary, it was designated as a DEG in the tumor. The final dataset included 5,097 tumors with 1,364,207 SGA events and 13,549,660 DEG events, **b.** A set of SGAs and a set of DEGs from an individual tumor as input for TCI modeling, **c.** The TCI algorithm infers the causal relationships between SGAs and DEGs for a given tumor *t* and output a tumor-specific causal model, **d.** A hypothetic model illustrates the results of TCI analysis. In this tumor, *SGA_SET_t_has* three SGAs plus the non-specific factor *A*_0_, and *DEG_SET*_*t*_ has six DEG variables. Each *E*_*i*_ must have exactly one arc into it, which represents having one cause among the variables in *SGA_SET_t_.* In this model, *E*_1_ is caused by *A*_o_;*E*_2_, *E*_3_, E4 are caused by *A*_*1*_; *E*_*5*_, *E*_6_ are caused by *A*_*3*_; *A*_*2*_ does not have any regulatory impact.

Specifically, we collectively evaluate the impact of different genomic alterations—in this study, we only consider SMs and/or SCNAs—affecting a gene as an SGA event, and we designate such an event using the gene name. The TCI model assumes that DEGs in an individual tumor are either caused by the SGAs in the tumor or by non-SGA factors related to tumor micro-environment (e.g., hypoxia). It further assumes that each DEG is likely regulated by one aberrant pathway in a tumor and such a pathway is likely perturbed by a single SGA, due to the well-known mutual exclusivity among SGAs perturbing a common pathway (32-34). As such, the TCI model constrains each DEG to be caused by one SGA in an individual tumor.

Given a collection of genomic data (denoted as D) from TCGA and the data from a new tumor *t,* in which an SGA event in a gene *h* is observed (denoted as *A*_*h*_) and a gene *i* is differentially expressed (denoted as *E*_*i*_), TCI evaluates the posterior probability of a model that the SGA event causes (drives) the DEG event, denoted as *A*_*h*_ → *E*_*i*_, in a Bayesian framework: *P*(*A_h_* → *E*_*i*_*|D*) ∝ *P*(*A*_*h*_ → *E*_*i*_)*P*(*D|A*_*h*_ → *E*_*i*_). The TCI framework involves the calculation of two components: a prior probability that *A*_*h*_ cause *E*_*i*_, i.e., *P*(*A_h_* → *E*_*i*_), and conditional probability (marginal likelihood) of data D given the model *P*(*D|A_h_* → *E*_*i*_). This setup allows us to integrate the assumptions of frequency-based framework and function-oriented framework.

Using an informative prior that express the possible biological impact of different genome alterations can help identify a correct model(35). We assessed the prior probability that *A*_*h*_ is likely a driver in a specific cancer type by adopting the assumption and approach of contemporary frequency-based driver framework (16), i. e., the more the SGAs in a gene is enriched in a cohort, the more likely it is a driver gene in an individual tumor. This allowed us to directly transform the results of the statistical test on the significance of mutations in a gene to a prior probability that *Ah* is a driver event in a tumor (see Methods).

We then evaluated the functional impact of an SGA *A*_*h*_ on expression of *i*^th^ gene *E*_*i*_, denoted as *A*_*h*_ → *E*_*i*_, using a tumor-specific marginal likelihood *P*(*D|A_h_ → E_i_).* The posterior probability *P*(*A_h_ → E_i_|D)* is derived and normalized by considering all SGA events in the tumor, so that the same edge *A*_*h*_ *→ E_i_* would have different posterior probabilities in different tumors because each tumor has a distinct *SGA*_*SET*_t_(Supplementary Method). In sum, TCI combines frequency-based approach and function-oriented framework to detect functions of candidate drivers in a tumor-specific fashion.

We applied TCI to analyze data from 5,097 tumors across 16 cancer types in TCGA (https://cancergenome.nih.gov/, Supplementary Table SI) to derive 5,097 tumor-specific models (a causal network model per tumor). We defined an SGA event in a tumor as an SGA with functional impact (SGA-FI) if it was predicted by TCI to causally regulate 5 or more DEGs in the tumor with an expected false discovery rate ∼ 10∼^7^, which is determined based a series of random simulation experiments. (Methods and Supplementary Figure SI).

We have identified a total of 634 genes that were called as SGA-FIs in more than 30 tumors with SGA-FI call rate^1^ of 25% or greater in our pan-cancer analysis (Methods and Supplementary Table S2). These SGA-FIs include the majority (87%) of the previously published drivers *(1,2),* as well as many novel candidate drivers. In addition to protein-coding genes, TCI also identified SGA events affecting microRNAs and intergenic non-protein-coding RNAs (e.g., *M1R31HG(36,37), MIR30B(38),* and *PVT1(39’))* as SGA-FIs, (Supplementary Table S2).

We further identified target DEGs for 634 significant SGA-FIs. To minimize false discovery, we required that a target DEG of an SGA-FI be regulated by the corresponding SGA-FI in at least 50 tumors or 20% of all tumors in which the SGA was called as an SGA-FI. Since it is statistically difficult to evaluate whether the causal relationship between an SGA-FI and its predicted target DEG within an individual tumor is valid, we adopted a “pan-cancer” analysis approach to determine whether each predicted SGA → DEG causal relationship is conserved across tumors in different cancer types. To minimize the confounding effect produced by tissue-specific SGAs and tumor-specific DEGs during pan-cancer analysis, we also performed TCI analysis on tumors from each cancer type separately, and we only retain SGA-FIs detected in at least two cancer types in tissue-specific analysis (Supplementary Table S2). We then set out to assess whether the inferred causal relationships are supported by existing knowledge and experimental studies. Finally, we performed preliminary laboratory experiments on selected SGA-FIs to evaluate the causal relationships between novel candidate drivers and their target DEGs predicted by TCI.

### The landscape of causative SGAs identified by TCI

We compared the distribution of the number of SGAs and SGA-FIs per tumor across cancer types (Figure 2a - 2b). The average number of SGAs per tumor across cancer types was 268, whereas the average number of SGA-FIs identified by TCI was approximately 34 per tumor. Interestingly, TCI designated all SGAs with very high alteration frequency (perturbed in more than 500 tumors, or > 10%) as SGA-FIs (Figure 2c). One immediate concern for TCI is that it could call certain long genes, such as *TTN* and *MUC16,* as SGA-FIs solely due to their high genomic alteration rate. This concern was addressed by adopting statistical test results from MutSigCV analysis, which specifically addressed the biased mutation rate introduced by lengths and chromosome locations of genes. We transformed such knowledge as the prior probability *P*(*A_h_ → E_t_)* in our analysis, and as such the prior probabilities for certain long genes, e.g., *TTN* and *MUC16,* were several orders of magnitude lower than other frequently altered well-known drivers (Figure 2d). This indicates that the strength of statistical relationships between SGAs in these genes and their target DEGs, reflected as the marginal likelihood *P*(*D|A_h_*→ *E*_*i*_), must be sufficiently high to overcome the low prior probabilities of these genes to be designated as regulators for DEGs. Many SGA-FIs with an alteration frequency ranging from 30 to 500 tumors (0.5 - 10%) are dispersed among other SGAs with similar protein lengths and alteration rates (Figure 2c). Since genes with similar protein length and alteration rate usually have similar prior probabilities of being drivers, TCI differentiated SGA-FIs from others based mainly on the difference in marginal P(D *|A_h_* → *E*_*t*_) associated with an SGA and its target DEGs. These results indicate that the function-oriented nature of TCI plays a significant role in detecting SGA-FIs.

**Figure 2.**
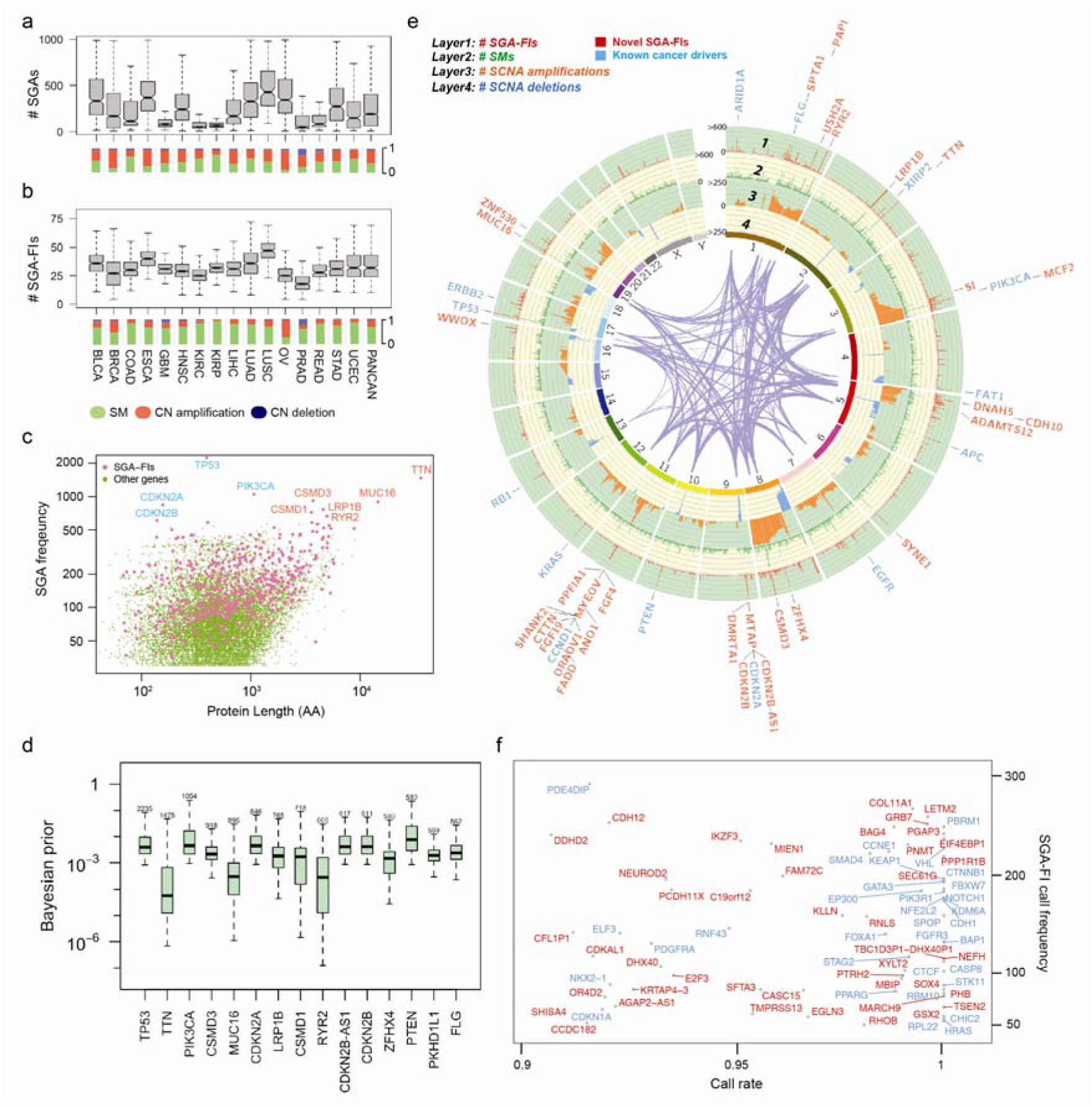
The landscape of SGAs and SGA-FIs. **a & b.** The distributions of SGAs per tumor and SGA-FIs per tumor of different cancer types. Beneath the bar box plots, the distributions of different types of SGAs (SM, copy number amplification, anddeletion) are shown, **c.** Identification of SGA-FIs is independent of the alteration frequency or protein length. Pink dots indicate SGA-FIs, and green dots represent SGAs that were not designated as SGA-FIs. A few commonly altered genes are indicated by their gene names, where genes labeled with blue font are well-known drivers, and those labeled with orange font are novel candidate driver, **d.** Tumor specific Bayesian prior distributions for top 15 most frequent SGAs. The number above each box represents number of tumors that the corresponding SGA appears in. **e.** A Circos plot shows SGA events and SGA-FI calls along the chromosomes. Different types of SGA evetns (SM, copy number amplification, deletion) were shown in the track 2, 3, and 4 respective. The track 1 shows the frequency of an SGA be called as SGA-FIs. The genes names denote the top 62 SGA-FIs (some are SGA units) that were called in over 300 tumors with a call rate > 0.8. Genes labeled with blue font are known drivers from two TCGA reports, and orange ones are novel candidate drivers, **f.** SGA-FIs that were called in less than 300 tumors and with a call rate >0.9 are shown in this frequency-vs-call rate plot. Similarly, genes labeled with blue font are known drivers from TCGA studies, and orange ones are novel candidate drivers.

We illustrated the landscape of common SGA events (Figure 2e) using a Circos plot (http://circos,ca/). and highlighted 44 SGA-FIs identified by TCI in more than 300 tumors (> 6% of the tumors) with a call rate (fraction of SGA instances affecting a gene being called as an SGA-FI event) greater than 0.8. The plot illustrates the integrative approach of TCI, which combines different types of SGA events in a gene and detects their function impact. For example, TCI combined mutation and deletion events in *LRP1B* (at 1 to 2 o’clock position on the plot) to detect common functional impact of these SGA events (see later section), whereas calling SGA-FI events for *ERBB2* (Her2) is mostly associated with amplification of the gene. TCI also designated many relatively low-frequency SGAs as SGA-FIs (in ∼ 30 tumors or ∼ 0.5%) with high call rates (> 0.9) (Figure 2f). Of interest, besides identifying well-known cancer drivers, e.g., *TP53, PIK3CA, PTEN, KRAS,* and *CDKN2A*) as SGA-FIs, TCI also designated as SGA-FIs some very frequently altered genes, e.g., *TTN, CSMD3, MUC16, LRP1B,* and *ZFHX4,* whose roles in cancer development remain controversial. These genes are excluded from driver gene lists when assessed by mutation-centered and frequency-based methods (2,16), but other computational and experimental studies (40-42) suggest that some of them are likely cancer drivers.

### Combining SM and SCNA enhances detection of the functional impact of genes affected by SGAs

A cancer driver gene is often perturbed by different types of SGA events that exert common functional impact. For example, an oncogene, e.g., *PIK3CA,* is usually affected by activating mutations or copy number amplifications, whereas a tumor suppressor, e.g., *PTEN,* is usually affected by inactivating mutations or copy number deletions. An SCNA event (amplification or deletion of a chromosome fragment) in a tumor often encloses many genes, making it a challenge to distinguish the functional impact of genes within a SCNA fragment.

TCI addresses this problem by integrating both SM and SCNA data, which can create variances in overall SGA events among genes within a SCNA fragment. When combined with SM data, *PIK3CA* clearly has a higher combined alteration rate than its neighbor genes in the same DNA region with very similar amplification rate, i.e. cytoband 3q26. While its neighbor genes share almost identical copy number amplification profile across all tumors, the alteration profile of *PIK3CA* are significantly different if both SM and SCNA data are considered (Figure 3a). When calculating whether amplification of *PIK3CA* is causally responsible for a DEG observed in a tumor, TCI uses the statistics collected from all tumors with *PIK3CA* alterations, including both CN amplification and SM, to compute the marginal likelihood and evaluate whether the causal relationship between *PIK3CA* amplification and the DEG is preserved in the tumor. As such, the algorithm is able to differentiate the functional impact of *PIK3CA* amplification from that of other coamplified genes. We noted that many genes were affected by both SMs and SCNAs patterns, including *CSMD3* and *ZFHX4* (Figure 3a), enabling TCI to detect the functional impact of these SCNA events.

**Figure 3.**
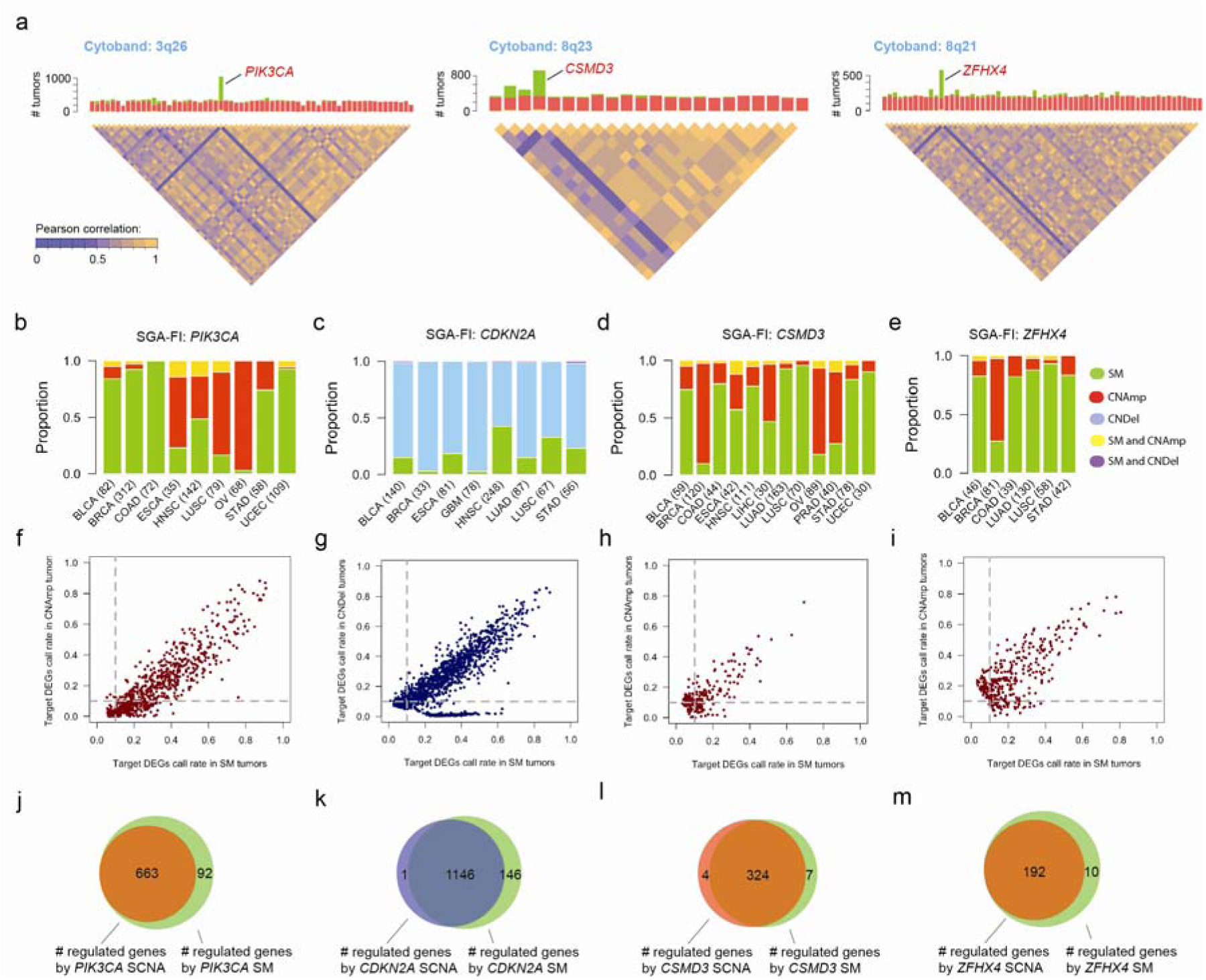
SM and SCNA perturbing a gene exert common functional impact, **a.** Combining SM and SCNA data drisrupts the correlation structure among genes enclosed in a common SCNA fragments. The chromosome cytobands enclosing three example genes (*PIK3CA, CSMD3, and ZFHX4*) are shown. The bar charts show the frequency of SCNA (red, standing for amplificaton) and SM (green). The disequilibrium plots beneath the bar charts depict the correlationship among genes within a cytoband. **b-e.** The SGA patterns, i.e. SM and CN amplification/deletion, across different cancer types for *P1K3CA, CDKN2A, CSMD3* and *ZFHX4,* f-i. SGA-FI target DEGs call rate in SM tumors and CN amplification/deletion for *PIK3CA, CDKN2A, CSMD3* and *ZFHX4.* **j-m.** Venn diagrams illustring the relationships of DEGs caused by CN amplication/deletion and SM for *P1K3CA, CDKN2A, CSMD3* and *ZFHX4.*

By combining both SM and SCNA data, TCI is able to identify common functional impact of distinct types of SGA events affecting the same gene across different tumors and cancer types. For example, *PIK3CA* is often perturbed by either SMs or CN amplifications (Figure 3 b) although prevalence of each type is different in different cancer types. In breast cancers, *PIK3CA* is commonly altered by SMs; in ovarian cancers, it is more often affected by CN amplification; in head and neck squamous carcinoma, it is almost equally altered by SMs and CN amplification. As a well-known cancer driver in many cancer types, it is expected that amplification and mutations of *PIK3CA* should share a common functional impact, i.e., causally regulating a common set of DEGs.

Taking advantage of the tumor-specific inference capability of TCI analysis, we identified the target DEGs regulated by each SGA event affecting *PIK3CA* (either SM or SCNA) in individual tumors. DEGs predicted to be caused by either *PIK3CA* SM or CN amplification have very similar positive predictive values (PPV) with respect to SGA events in *PIK3CA.* PPV is calculated as the ratio of number of tumors in which a DEG is designated as target of an SGA-FI such as *PIK3CA* over all tumors in which *PIK3CA* is called as an SGA-FI (Figure 3f and Methods). The results indicate that perturbation of *PIK3CA* by both SM and CN amplification have very similar functional impact on gene expression changes. We then examined whether target DEGs caused by *PIK3CA* SM overlap with those caused by CN amplification, and indeed the DEG members of the two list significantly overlapped (Figures 3j). Thus, TCI detected the shared functional impact of distinct types of SGAs perturbing *PIK3CA* across different cancer types. Similar results were obtained for other 249 SGA-FIs (Supplementary Table S3) that were commonly perturbed by both SMs and SCNAs (with each type accounting for > 20% of instances for each SGA-FI), including *CDKN2A, CSMD3* and *ZFHX4* (Figures 3).

### Causal relationships inferred by TCI are statistically robust

To evaluate validity of the results by TCI, we first examined whether the causal relationships reported by TCI reflect true statistical relationships between SGA and DEG events rather than random noise in data. We generated a series of random datasets using the TCGA data, in which the DEG status of each gene expression variable was permuted among the tumors, while the SGA status in each tumor remained as reported by TCGA. After permutation, the statistical relationships between SGAs and DEGs are expected to be random. We then applied TCI to these random datasets and compared the posterior probabilities of the most probable causal edges for each DEG derived using real and permuted data. The results (Figure 4a) show TCI were able to differentiate true statistical relationships between SGAs and DEGs from random ones in that it assigned higher posterior probabilities to candidate edges obtained from real data (red lines) than those obtained from random data (blue lines). As expected, a large number of derived causal edges from well-known cancer drivers (e.g., *TP53* and *PIK3CA*) were assigned high posterior probabilities. Interestingly, the results also show many causal edges from other common SGA-FIs *(TTN, CSMD3, MUC16,* and *ZFHX4*) to DEGs were also assigned higher posterior probabilities than would be expected by random chance, indicating that perturbing these genes had significant impact on transcriptomics of the tumors (Figure 4a and Supplementary Figure S2a). The function-oriented nature of TCI is reflected by observations that there are certain SGAs with a high alteration frequency (occurring in close to 10% of tumors) were not designated as SGA-FIs by TCI. For example, *WASHC5* has SGA events in 457 tumors but few of these SGA events were assigned with high posterior probabilities, similar results were observed for *TBC1D31* (424 tumors) and *ADGRB1* (420 tumors) (Supplementary Figure S2b).

**Figure 4.**
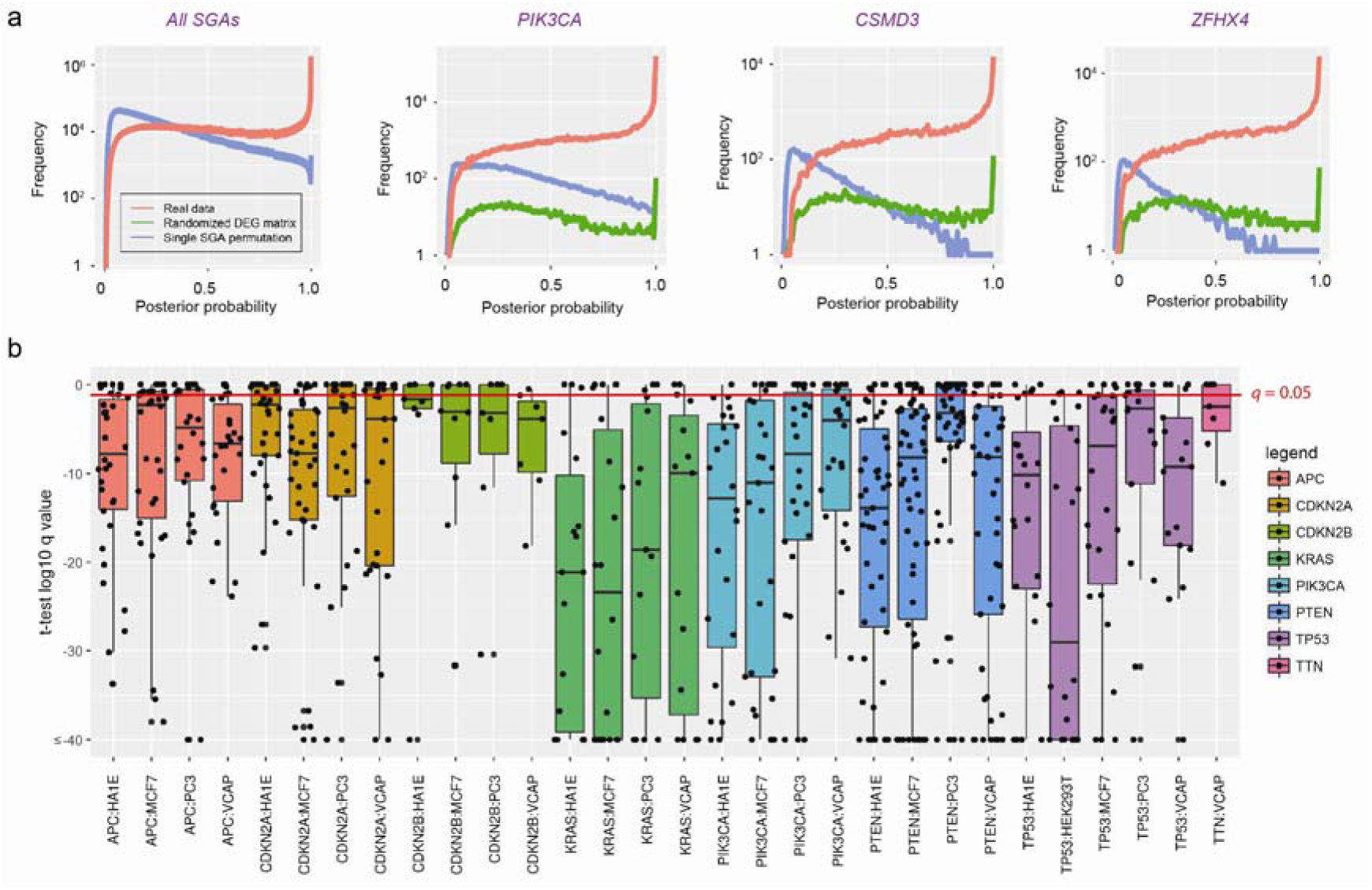
Statistical and experimental Evaluation of TCI predictions, **a.** The causal relationship inferred by TCI is statistically sound. Plots in this panel show the probability density distribution of the highest posterior probabilities assigned to each DEG in TCGA dataset, when the TCI algorithms was applied to real data (red) and two random datasets, in which DEGs permutated across all tumors (blue) and the corresponding SGA permutated across all tumors (green). The panel on the left shows the results for the posterior probabilities for all most probable candidate edges in whole dataset; rest of the plots show the distributions of posterior probaiblities of most probable edges pointing from 3 specific SGAs to predicted target DEGs. **b.** Boxplots of q-values associated with predicted target DEGs for 8 SGA-FIs in different LINCS cell line after perturbation experiments. Each black dot represent a *q*-value associated with a target DEG of an SGA-FI, when expression value was assessed with *t*-test contrasting before and after genetic manipulation of the corresponding SGA-FI gene.

To further exclude the possibility that TCI-reported causal relationships from a high-frequency SGA to DEGs were random associations due to their high alteration frequencies, we conducted another series of single-SGA-permutation experiments, in which the SGA events of a gene, e.g., *TTN,* were randomly permuted across all tumors to disrupt the statistical relationships between SGAs of this gene and DEGs, while the overall frequency of the SGAs of the gene remains the same. We performed such single-SGA-permutation experiments for the 6 most commonly altered genes: *TP53, TTN, P1K3CA, CSMD3, MUC16,* and *ZFHX4.* The results (Figure 4a and Supplementary Figure S2b) also shows that when TCI analysis was applied to these permuted data (green lines), none of these 6 genes were designated as an SGA-FI according to our criteria. Taken together, these results indicate that TCI captures the true statistical relationships between SGA and DEG events in real data.

### Causal relationships inferred by TCI are biologically sensible

We further evaluated whether the TCI-inferred causal relationships between SGAs and DEGs agree with existing knowledge and experimental results. We compared the predicted causal relationships between *PIK3CA* and DEGs with experimental results from an independent study. Recently, Hart et al. (43) studied the functional impact of single mutation, H1047R, of *PIK3CA,* by knocking in the mutation into the MCF10 cell line and comparing transcriptomic profile between the wild type and the *PIK3CA^m047R^* isogenic cell lines. They identified 1,434 DEGs caused by introduction of the mutation. We compared the TCI-predicted *PIK3CA* target DEGs with that from their study, and 12 out of 92 TCI-predicted *P1K3CA* SM driving DEGs overlaps with the experiment-derived DEG set (hypergeometric test *p* = 0.01).

Since RB1 protein regulates the function of transcription factor E2F1(44), it is expected that E2F1-regulated genes should be enriched among the *RBI*-targeted DEGs predicted by TCI. We used the PASTAA program(45) (trap.molgen.mpg.de/PASTAA.htm) to search for motif binding sites in the promoters of the 237 DEGs that TCI predicted to be regulated by *RB1,* and it found that *E2F1, E2F2,* and *DP-1* were the three top transcription factors for these genes (*p <10^-6^*)

In addition to verifying experimental evidence for individual SGA-FIs and their target DEGs, we took advantage of large-scale perturbation experiments carried by the Library of Integrated Network-Based Cellular Signatures (LINCS) project (46) to systematically evaluate predicted causal relationships between SGA-FIs and their predicted target DEGs. The LINCS project performed systematic gene-manipulation (knockdown and overexpression) experiments using small interfering RNAs targeting over 4,000 genes in multiple cell lines, and cellular responses were measured as expression changes in 978 landmark genes (using a technology referred to as L1000 assay). We selected 8 most frequent SGA-FIs that were experimentally manipulated in LINCS project and performed *t*-test on the expression values of all L1000 genes, contrasting perturbation experiments and control condition in each cell line. We then examined the statistical significance of these genes and assess the false discovery (q-values) associated with the predicted target DEGs of each SGA. For each of the 8 SGA-FIs, the majority of predicted target DEGs were differentially expressed in multiple cell lines after experimental manipulation of SGA-FI gene (Figure 4b). We noted that certain target DEGs of an SGA have tissue-specific expression patterns, and we organized targeted DEGs according to tissue of origins and examined the percent of DEGs responding to manipulation of corresponding SGAs (Supplementary Table S4). Interestingly, we also found that *TTN* was perturbed in one cell line (VCAP), and 5 out of 7 predicted target DEGs responded to manipulation of *TTN.* Among them, 4 genes (*SPP1, STAT1, C5* and *GPER1*) are known to be functionally associated with development and/or progression of cancer (47-50). To sum, the causal relationships between SGAs and DEGs predicted by TCI can be validated from multiple perspectives, including existing knowledge regarding genes, targeted or systematic experiment results.

Finally, we experimentally examined whether experimental manipulations of *CSMD3* and *ZFHX4* expression affect oncogenic phenotypes. We identified two cancer cell lines, HGC27 and PC3, with *CSMD3* and *ZFHX4* amplification respectively, and we knocked down the expression of the two genes using siRNAs, followed by monitoring cellular phenotypes. Our results showed that knocking down *CSMD3* and *ZFHX4* in the respective cell lines significantly attenuated cell proliferation (viability) and migration (Figure 5a - d). In addition, knockdown of *ZFHX4* induced apoptosis (Figure 5e). These results indicate that these genes are involved in maintaining the cancer-related cellular phenotypes in these cell lines.

**Figure 5.**
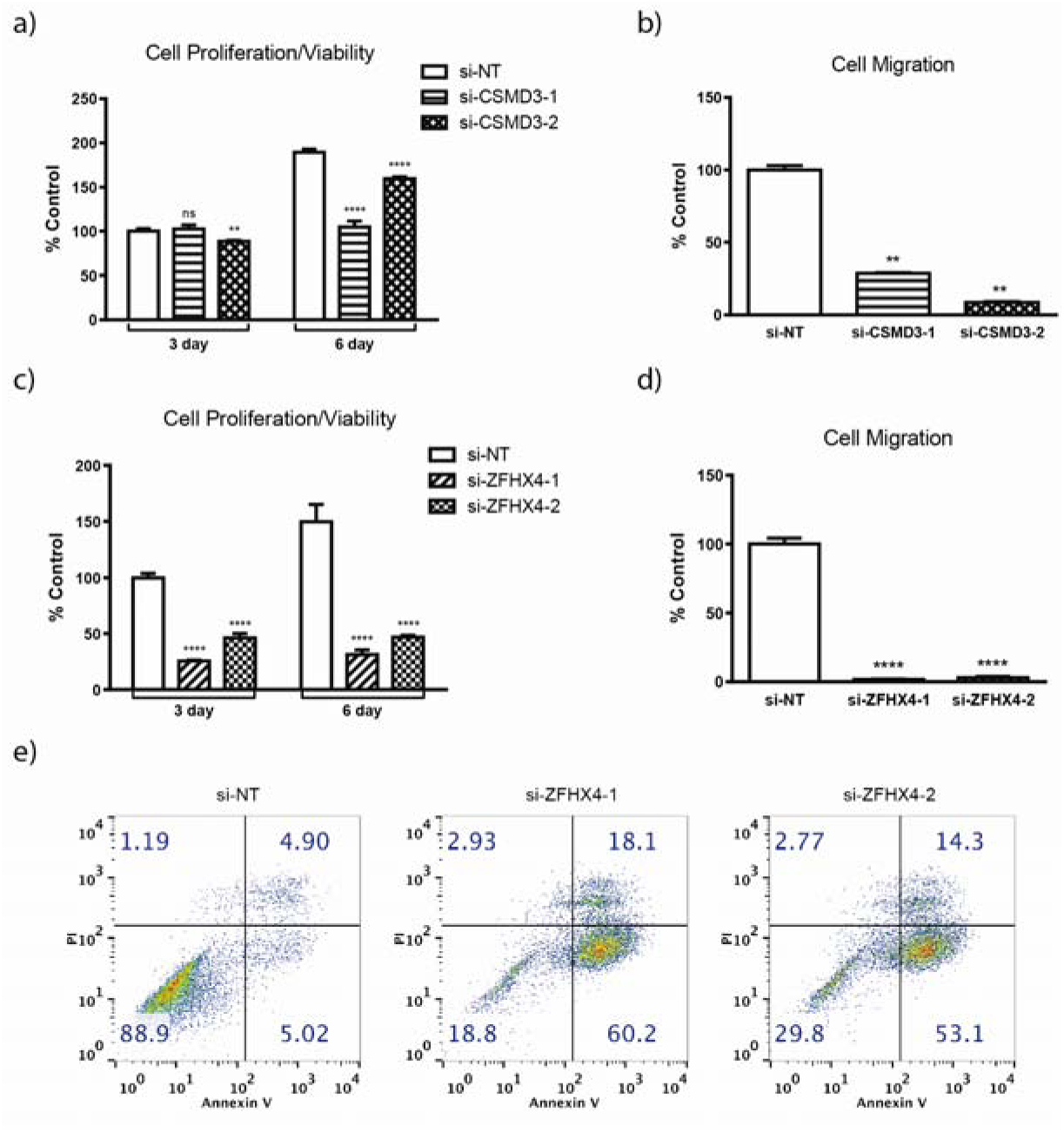
Cell biology evaluation of oncogenic properties of *CSMD3* and *ZHFX4.* **a-b.** The impact of knocking down *CSMD3* and *ZFHX4* on cell proliferation, **c-d.** The impact of knocking down *CSMD3* and *ZFHX4* on cell migration, **e.** Impact of *ZFHX4* knockdown on apoptosis in PC3 cell line measured by Annexin V and propidium iodide (PI) staining.

### SGA-FIs regulate genes involved in well-known oncogenic processes

To gain a better view of functional impacts of SGA-FIs in cancer development, we further examined their impact on 1,855 genes from 17 cancer-related “hallmark” gene sets from the MSigDB (http://software.broadinstitute.org/gsea/msigdb/index.jsp). On average, 374 cancer hallmark genes are found to be differentially expressed in a tumor. TCI found 96 SGA-FIs that are predicted to regulate members of these 1,855 hallmark genes, in other words, these results illustrate the impact of an SGA on cancer hallmark process. We listed the relationships between 96 SGA-FIs with respect to the 17 cancer hallmark processes to identify the target DEGs for each of 96 SGA-FIs (Supplementary Table S5). The relationships between top 45 SGA-FIs with largest number of target DEGs with respect to the hallmark processes are shown in (Figure 6a). For example, *CTNNB1* is known as the top regulator of WNT pathway and it is predicted to cause 14% DEGs in HALLMARK_WNT_BETA_CATENIN_SIGNALING pathway (51); *RB1* regulates 15% of the genes in HALLMARK_E2F_TARGETS (52); *TP53* regulates genes involved in apoptosis and in a broad assortment of functions across many other oncogenic pathways (53,54); Our analysis also suggests that *CDKN2A* plays an important role in the epithelial-mesenchymal transition (EMT) process, which agrees with previous studies (55).

**Figure 6.**
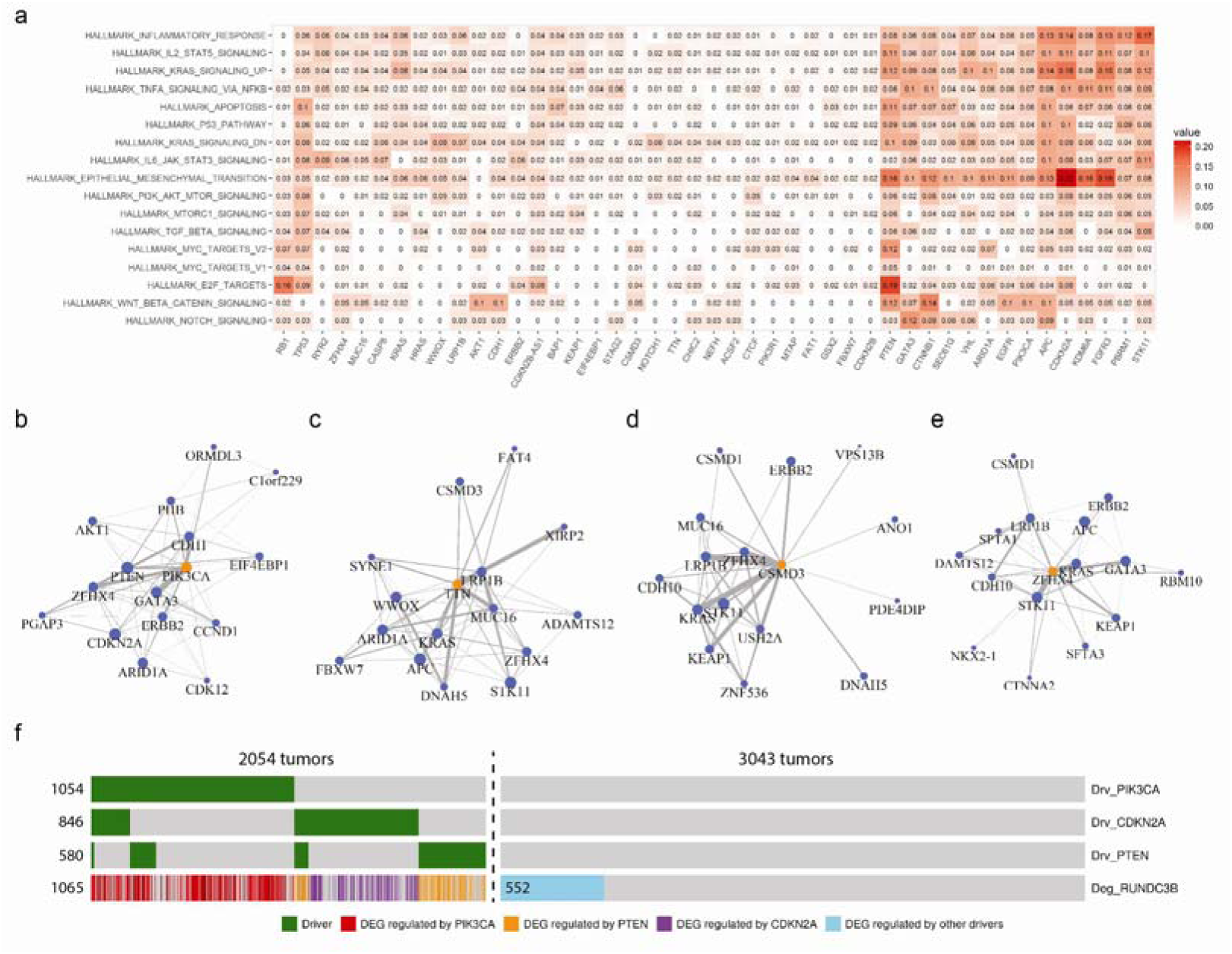
Detection of functional impact of SGA-FIs reveals functional connections among SGA-FIs. **a.** Top 45 SGAs-FIs (regulating the largest number of DEGs) and their relationships with 17 cancer hallmark gene sets. The value represents the percentage of genes in a gene set that is covered by the target DEGs of each SGA-FI. **b-e.** Top 15 SGA-FIs that share the most significant overlapping target DEGswith *P1K3CA, TTN, CSMD3,* and *ZFHX4.* An edge between a pair of SGA-FI indicate that they share significantly overlapping target DEG sets, and the thickness of the line is proportional to negative log of the p-values of overlapping target DEG sets. **e.** The “oncoprint” illustrating the causal relationships between a DEG, *RUNDC3B,* and its 3 main drivers, *P1K3CA, CDKN2A,* and *PTEN.* Each column corresponds to a tumor; green bars show SGA events that cause *RUNDC3B* DEG in 3 SGA-FIs; the causal relationship is color-coded, which illustrate which SGA-FI is responsible for a DEG event in *RUNDC3B;* blue bar indicates that DEG events that were assigned to SGA-FIs other than the above 3 SGA-FIs; gray bars indicate a wild type genomic and transcriptomic status.

### TCI analyses reveal functional connections among SGA-FIs

The causal relationships between SGAs and DEGs revealed by TCI enable us to explore whether distinct SGAs in different tumors are in fact perturb a common signal, by examining if they share overlapping target DEGs. To this end, we evaluated all pair-wise intersections between target DEG sets of SGA-FIs to identify SGA pairs sharing significantly overlapping target DEGs (p <0.05 Fisher’s exact test, and *q <*0.05), and found 2669 such SGA-FI pairs (Supplementary Table S6). We then organized SGA-FIs that perturb common signals into a graph, in which an edge connecting a pair of SGA-FI nodes indicates significant overlap of their target DEGs. For example, the top 15 SGA-FIs (ranked according to the FDR *p* values of overlapping DEG sets) that share DEGs with *P1K3CA* include *PTEN, CDH1, ERBB2,* and *GAT A3,* which are known cancer drivers, and their connections agree with existing knowledge (Figure 6b) (56-58).

The capability of revealing functional connections among SGAs provides a means to evaluate whether a novel candidate driver shares functional impact with well-known drivers, which not only indicates whether the candidate driver is involved in oncogenic processes (and thus a candidate cancer driver gene) but also sheds light on which pathway it may be involved in. The top 15 SGA-FIs sharing common target DEGs with *TTN* include some well-known drivers including *APC, KRAS, STK11* and *SYNE1* (Figure 6c). Therefore, *TTN* might share similar functional impact with these known drivers. The top 15 SGA-FIs connected with *CSMD3* and *ZFHX4* (Figure 6d and Figure 6e) also form densely connected networks that include well-known cancer drivers, such as *KRAS, GATA3, KEAP1, ERBB2* and *STK11,* suggesting that alteration of *CSMD3* and *ZFHX4* perturb some of the same signaling pathways as these known drivers do. We found similar results for other common SGA-FIs, including *CDKN2A, PTEN, MUC16,* and *LRP1B* (Supplementary Figure S3).

Transcription of a gene is often regulated by a pathway, and it is expected that major driver SGAs of a DEG should include members of such a regulatory pathway. As an example, Figure 6f shows the SGA events that TCI designated as the cause of differential expression of *RUNDC3B* in different tumors, for which TCI analysis indicates that *PIK3CA* is the most common cause. Besides SGAs in *PIK3CA,* TCI inferred that SGAs in *CDKN2A, and PTEN* are two other major drivers of *RUNDC3B* DEG events. The results suggest that aberrations in PI3K pathway (as a result of SGAs perturbing *PIK3CA* and *PTEN)* is the main cause of these DEG events, and *CDKN2A* may act as a downstream regulator. It is also interesting to note that, in certain tumors, when both SGAs affecting *PIK3CA* and *PTEN* were present, TCI assigned *PTEN* as the most likely driver of *RUNDC3B,* instead of *PIK3CA,* even though the SGAs in the latter is more frequent. The results indicate that, although *PIK3CA* SGA events explains the overall DEG variance of *RUNDC3B* better than *PTEN,* the strength of statistical association between *PTEN* and some DEGs in certain tumors may be stronger than that of *PIK3CA,* and TCI can detect such statistical relationships.

### Tumor-specific causal inference reveals tumor-specific disease mechanisms

TCI analysis enables us to identify major SGAs that causally regulate molecular phenotypic changes (in our case, DEGs) in an individual tumor. In this way, TCI not only discovers potential drivers of an individual tumor but also suggests which oncogenic processes they may affect. In other words, TCI can reveal tumor-specific disease mechanisms, particularly when more oncogenic phenotypic data types become available.

TCI results enabled us to examine each tumor profiled by TCGA to identify the major candidate driver SGAs and their target DEGs. Further examining the DEGs involved in hallmark biological processes allows us to study which biological processes an SGA affects. As an example, Figure 7a shows the SGA-FIs and their target cancer processes for a tumor (TCGA-B1-A657) of Kidney Renal Papillary cell carcinoma (KIRP), where genes in 9 oncogenic hallmark process from MSigDB are significantly enriched among the DEGs, including the following pathways that are strongly regulated by one of more SGA-FIs: the Epithelial Mesenchymal Transition pathway, the KRAS signaling pathway, the TNFA signaling via NFKB pathway, and the IL2 STAT5 signaling. We also identified major SGA-FIs (according the number of DEGs regulated by them in the tumor) that affect these processes (Figure 7a). In this figure, a green arrow indicates that an SGA-FI regulates at least 10% of the genes in the corresponding signaling pathway. TCI identified 6 such SGA-FIs, including some well-known cancer drivers, such as *PTEN* and *NEFH,* and potential cancer drivers mentioned in recent studies, such as *TLK2(*59), *USP13(*60), and *P1M3(*61).

**Figure 7.**
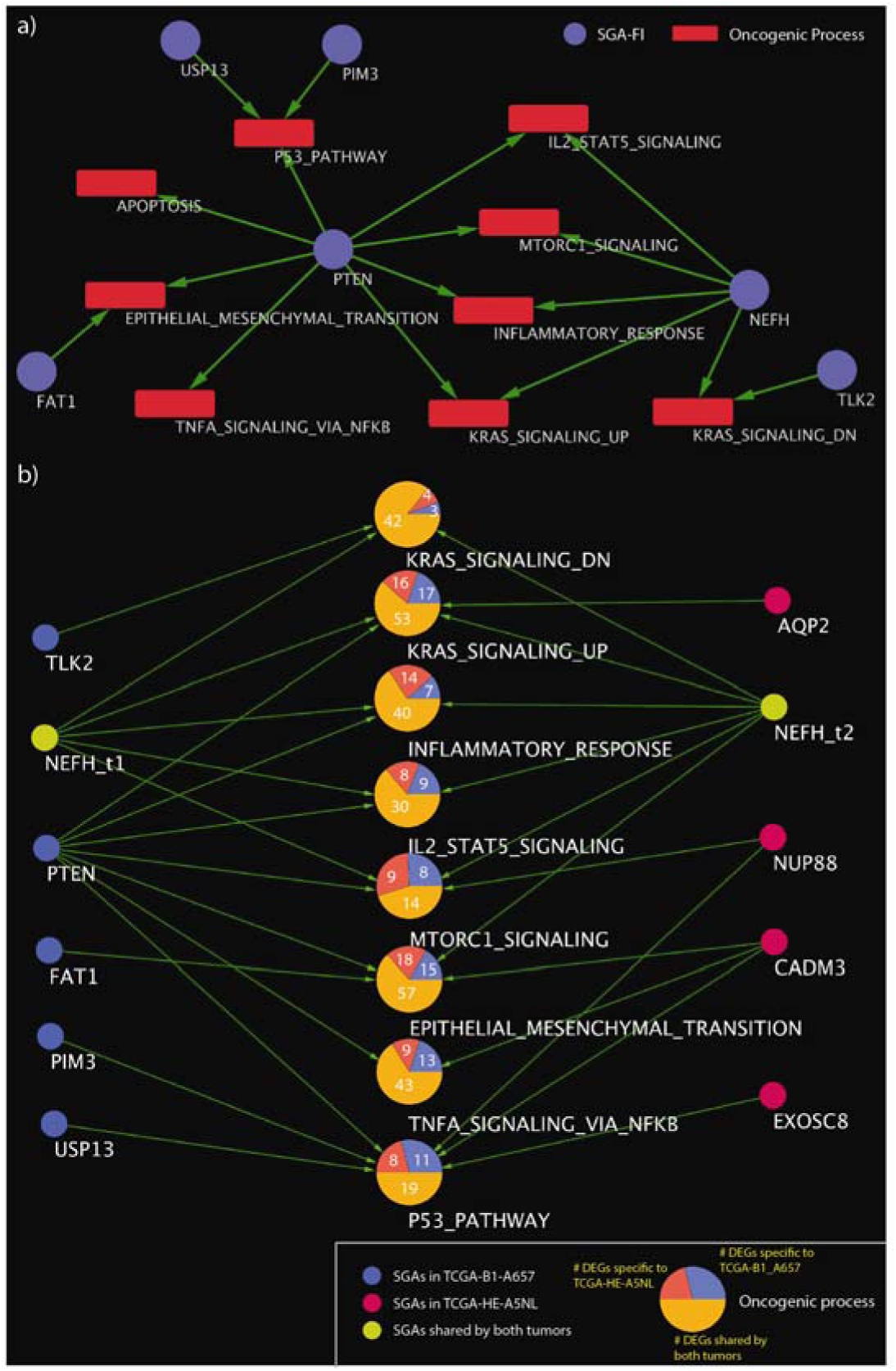
TCI reveals the SGA-FIs and their functional impacts at the individual tumor level, **a.** The bipartite graph illustrating major SGA-FIs and their regulated cancer processes for the tumor TCGA-B1-A657. Blue node represent SGA-FIs and red nodes (square) represent oncogenic processes. An green directed link indicates an SGA-FI regulates 10% or more DEGs in the cancer process, **b.** Same DEGs regulated by distinct SGA-FIs in different tumors. DEGs in cancer processes shared between tumor TCGA-B1-A657 and tumor TCGA-HE-A5NL are shown as pie-charts. Blue nodes denote SGA-FIs in tumor TCGA-B1-A657. Red nodes denote SGA-FIs in tumor TCGA-HE-A5NL. Yellow nodes, (i.e., *NEFH),* is shared by both tumors. Each large node in the middle represents an oncogenic process. The number in the blue area denotes the number of DEGs specific to TCGA-B1-A657. The number in the red area denotes the number of DEGs specific to TCGA-HE-A5NL. The number in the yellow area denotes the number of DEGs shared by both tumors. An green directed link indicates an SGA-FI regulates 10% or more DEGs in the cancer process.

SGAs cause cancer by perturbing cellular signaling pathways, and a pathway usually consists of multiple signaling proteins. Thus, it is possible that tumors having very distinct SGA profiles may in fact share very similar patterns of pathway perturbation, thus sharing similar gene expression profiles. We further identified another KIRP tumor (TCGA-HE-A5NL), which shares a similar overall DEG profile as does TCGA-B1-A657 (Figure 7b). These two tumors shared 281 DEGs related to the aforementioned oncogenic processes, and many DEGs in each oncogenic process were shared by the two tumors. However, each of these two tumors also had its unique SGA set, such as 57 SGAs in TCGA-B1-A657, 65 in TCGA-HE-A5NL, and only 2 common SGAs (*CADM3* and *NEFH).* TCI discovered similar target DEGs for *NEFH* in both tumors. Although many DEG members in each oncogenic process were shared, different SGA-FIs were designated as their candidate drivers. The above results illustrate that TCI is able to suggest disease mechanisms of individual tumors, and such information can be further analyzed to suggest tumors sharing common disease mechanisms.

## Discussion

In this report, we present the TCI algorithm, which concentrates on addressing a fundamental question in discovering cancer-driving genes: whether perturbation of a gene (considering different perturbation mechanisms) is causally responsible for certain molecular/cellular phenotypes (considering different phenotypic measurements) relevant to cancer development in a tumor. Thus, TCI provides a principled statistical framework for identifying causative SGAs and understanding their functional impact on oncogenic processes of an individual tumor.

TCI is also a unifying framework for combining the statistics of all types of SGA events affecting a gene to assess whether perturbation of the gene causally lead to one or more molecular/cellular phenotypes. This causality-centered framework circumvents the need of separately assessing whether mutations, or SCNAs, or other SGA events in a gene are over-enriched in a cancer population by conventional approaches, which would require dealing with unconformable measurements and baseline models associated with each type of SGA. Integrating diverse types of SGA events is statistically sensible which increases the statistical power for assessing the functional impact of perturbing a candidate driver gene. It is also biologically sensible that a driver gene is often perturbed by different types of SGA events leading to common functional impact. The fact that a gene is often perturbed by different types of SGAs leading to common phenotypic changes provides strong evidence support that the gene is a candidate driver because its functional impact is positively selected in cancer.

Our analyses of TCGA data revealed the functional impact of many well-known as well as a large number of novel SGA-FIs with a wide range of prevalence in tumors ranging from 1% to more than 10%. These results serve as a catalogue of major SGA events that potentially contribute to cancer development. Discovery of novel candidate drivers also provide potential targets for developing new anti-cancer drugs. By revealing the functional impact of candidate drivers (e.g., a signature of DEGs), TCI results can be utilized to identify SGAs sharing similar functional impact and to discover cancer pathways *de novo* or to map novel candidate drivers to known pathways.

Interestingly, TCI revealed functional impact of certain SGAs with very high alteration frequencies, such as *TTN, CSMD3, MUC16, RYR2, LRP1B,* and *ZFHX4,* whose roles in cancer development remains controversial. There are studies indicating that their high mutation rates are likely due to heterogeneous mutation rates at different chromosome locations (2,16). TCI analysis provides a new perspective to examine the role of these genes: assessing whether perturbations (considering all SGA events) in these genes are causally responsible for molecular and cellular phenotype changes. Instead of concentrating on assessing whether its frequency is above random chance, TCI evaluate the functional impact of an altered gene that determines whether it contributes to (drives) cancer development. Our results suggest that perturbing these genes, either by genome alterations, such as SM and/or SCNA, or by experimental manipulations, bears significant impact on molecular and cellular changes in both tumors and cell lines. Therefore, our results motivate further investigation of an alternative hypothesis for high overall alteration rates of these genes in cancer: perturbation of these genes, despite of different ways, leads to functional changes that provide oncogenic advantages. The results suggests that utilizing diverse types of SGA events in these genes is in fact a result of positive selection.

TCI is a special case of the general instance-based causal inference framework (62,63) that can be broadly used to delineate causal relationships between genomic variance and phenotype changes at the level of individuals which can be a single cell or an individual patient. By studying tumors in TCGA dataset, TCI analysis sheds light on the disease mechanisms of each individual. Further exploring the commonality and differences in disease mechanisms for individual tumors in the population will significantly help us better understand cancer biology in general. More importantly, understanding the disease mechanism of each tumor lays a solid foundation for guiding personalized therapies and advancing precision oncology.

## Acknowledgments

The authors acknowledge editorial assistance provided by Michelle Kienholz and technical assistance by Fan Yu and Soumya Luthra. The authors would like to thank Drs. Clark Glymour, Peter Spirtes, and Josh Stuart for discussions and suggestions. Research reported in this publication was supported by grant U54HG008540 awarded by the National Human Genome Research Institute through funds provided by the trans-NIH Big Data to Knowledge (BD2K) initiative (www.bd2k.nih.gov). Funding also came from R01LM012011 and R00LM011673 awarded by the National Library of Medicine, and from Grant #4100070287 awarded by the Pennsylvania Department of Health and from Grant # PC 150190 awarded by the Department of Defense. The content is solely the responsibility of the authors and does not necessarily represent the official views of the National Institutes of Health or the Pennsylvania Department of Health or the Department of Defense.

## Methods

### SGA data collection and preprocessing

We obtained SM data for 16 cancer types directly from the TCGA portal f https://tcga-data.nci.nih.gov/tcga/dataAccessMatrix.htm) (accessed in October 2014). We considered all the non-synonymous mutation events of all genes and considered the mutation events at the gene level, where a mutated gene is defined as one that contains one or more non-synonymous mutations or indels.

SCNA data were obtained from the Firehose browser of the Broad Institute (http://gdac.broadinstitute.org/). TCGA network employed GISTIC 2.0(27) to process SCNA data, which discretized the gene SCNA into 5 different levels: homozygous deletion, single copy deletion, diploid normal copy, low copy number amplification, and high copy number amplification. We only included genes with homozygous deletion or high copy number amplification for further analysis. We further screened out the genes with inconsistent copy number alteration across tumors in a given cancer type (i.e., gene was perturbed by both copy number amplification and deletion events in the same cancer type and both types of events occurred > 25% of tumors).

We combined preprocessed SM data and SCNA data as SGA data, such that a gene in a given tumor was designated as altered if it was affected by either an SM event and/or an SCNA event.

### DEG data collection and preprocessing

Gene expression data were preprocessed and obtained from the Firehose browser of the Broad Institute. We used RNASeqV2 for cancer types with expression measurements in normal tissues. For cancer types without RNASeqV2 measurements in normal cells (i.e., glioblastoma multiforme and ovarian cancer), we used microarray data to identify DEGs. We determined whether a gene is differentially expressed by comparing the gene expression in the tumor cell against that in the corresponding tissue-specific normal cells. For a given cancer type, assuming the expression of each gene (log 2 based) follows Gaussian distribution in normal cells, we calculated the *p* values of each gene in a tumor, which estimated how significantly different the gene expression in tumor was from that in normal cells. If the *p* value was equal or smaller than 0.005 to either side, the gene was considered as differentially expressed in the corresponding tumor. Furthermore, if a DEG was associated with the SCNA event affecting it, we removed it from the DEG list of the tumor. We also removed tissue-specific DEGs if they were highly correlated with cancer types or tissue origin (i.e., Pearson correlation coefficient larger than 0.9). We thus identified the DEGs for each tumor and created a tumor-gene binary matrix where 1 represents expression change, and 0 represents no expression change.

### Identification of SGA-FIs

Causal edges from different SGAs have different posterior probabilities, as expected. To standardize how to interpret the significance of a posterior probability for a causal edge *P*_*e*_, we designed a statistical test based random permutation experiments. We generated a series of permuted datasets using the TCGA data, in which the DEG values were permuted among the tumors of a common tissue of origin, while the SGA status in each tumor remained as reported by TCGA. This permutation operation disrupts the statistical relationships between SGAs and DEGs while retaining the tissue-specific patterns of SGAs and DEGs. We applied TCI algorithm to permuted data to calculate posterior probabilities of edges emitting from each SGA in random data. We then determined the probability that an edge from an SGA could be assigned with a given *P*_*e*_ or higher in data from permutation experiments (i.e., the *p* value to the edge with a given *P*_*e*_).

The *p* value in this setting is also the expected rate of false discovery of an SGA as the cause of a DEG by random chance. We utilize this property to control the false discovery rate when identifying SGA-FIs in a tumor. We designated an SGA event in a tumor as an SGA-FI if it has 5 or more causal edges to DEGs that are each assigned a *p*-value <0.05. The overall false discovery rate of the joint causal relationships between an SGA to 5 or more target DEGs is smaller than 10^-7^. The Supplementary Figure S1 shows that at this threshold, none of SGA was assigned as SGA-FI by random chance.

### Cell culture and siRNA transfection

HGC27 (Sigma-Aldrich) and PC-3 (ATCC) cells were cultured according to the manufacturer’s recommendations. The non-targeting and the *CSMD3* and *ZFHX4* siRNAs were obtained from OriGene (Rockville, MD). The siRNA sequences are as follow: si-CSMD3-l, GGUAUAUUACGAAGAAUUGCAGAGT; si-CSMD3-2, ACAAAUGGAGGAAUACUAACAACAG; si-ZFHX4-1, CGAUGCUUCAGAAACAAAGGAAGAC; si-ZFHX4-2, GGAACGACAGAGAAAUAAAGAUUCA. The siRNAs were transfected into cells using DharmaFECT transfection reagents for 48 hrs according to the manufacturer’s instructions.

### Cell proliferation and viability assays

Cell proliferation/viability was assayed by CCK-8 assay (Dojindo Laboratories, Kumamoto, Japan). Briefly, HGC27 and PC3 cells were plated at a density of 3 × 10^3^ cells/well in 96-well plates. After siRNA transfection for 3 or 6 days, CCK-8 solution containing a highly water-soluble tetrazolium salt WST-8 [2-(2-methoxy-4-nitrophenyl)-3-(4-nitrophenyl)-5-(2,4-disulfophenyl)-2*H*-tetrazolium, monosodium salt] was added to cells in each well, followed by incubation for 1-4 h. Cell viability was determined by measuring the O.D. at 450 nm. Percent over control was calculated as a measure of cell viability.

### Trans well migration assay

Cell migration was measured using 24-well transwell chambers with 8 μm pore polycarbonate membranes (Corning, Corning, NY). SiRNA-transfected cells were seeded at a density of 7.5 × 10^4^ cells/ml to the upper chamber of the transwell chambers in 0.5 ml growth media with 0.1% FBS. The lower chamber contained 0.9 ml of growth medium with 20% FBS as chemoattractant media. After 20 hrs of culture, the cells in the upper chamber that did not migrate were gently wiped away with a cotton swab, the cells that had moved to the lower surface of the membrane were stained with crystal violet and counted from five random fields under a light microscope.

### Apoptotic assay

Apoptosis was assessed by flow cytometry analysis of annexin V and propidium iodide (PI) double stained cells using Vybrant Apoptosis Assay Kit (Thermo Fisher Scientific, Carlsbad, CA). Briefly, the cells after washing with PBS were incubated in annexin V/PI labeling solution at room temperature for 10 min, then analyzed in the BD FACSCalibur flow cytometer (Becton, Dickinson and Company, Franklin Lakes, NJ).

## Supplementary table legends

Supplementary Table S2.1. TCI predicted 634 Candidate SGA-Fis and their target DEGs

Supplementary Table S2.2. Cancer type distribution of 634 Candidate SGA-FIs

Supplementary Table S3. SGA-FIs that are commonly altered by both SM and SCNA

Supplementary Table S4. Number of target L1000 genes for 8 most frequent SGA-FIs that are differentially expressed in different tissue types

Supplementary Table S5. Percentage of genes involved in the cancer Hallmark processes regulated by SGA-FIs

Supplementary Table S6. SGA-FI pairs sharing common target genes

**Supplementary Figure S1.**
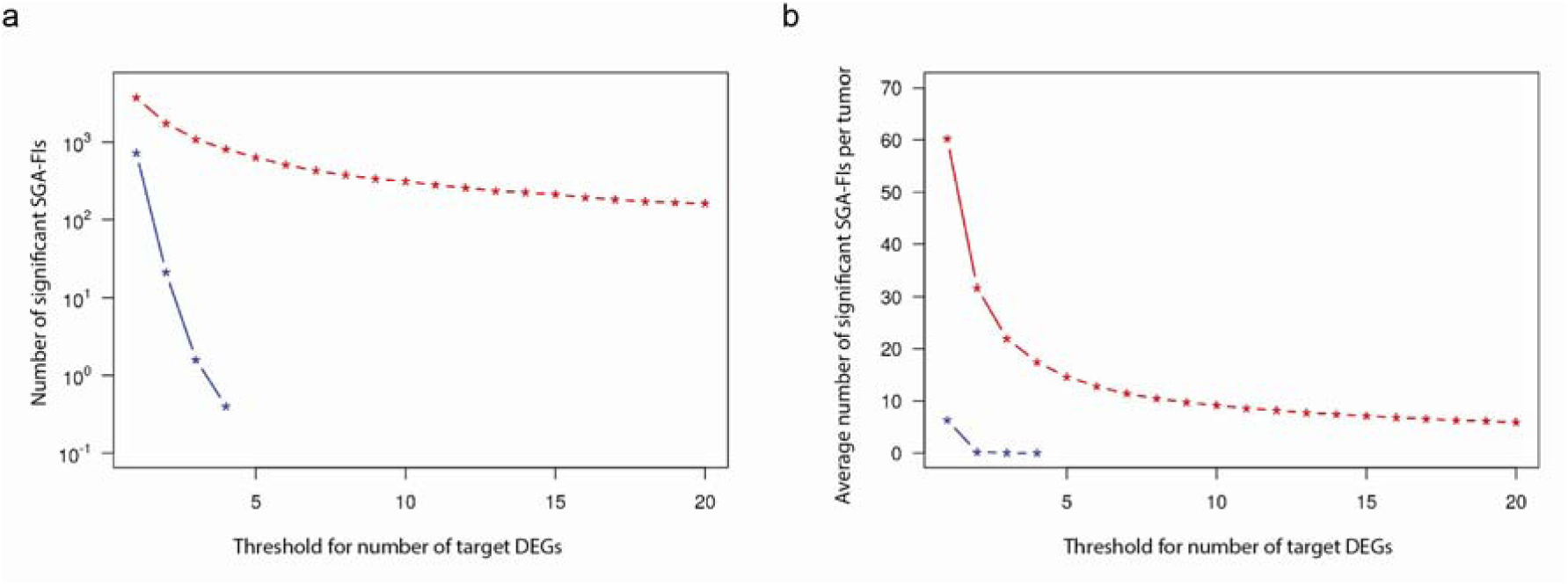
Comparison of causal analysis results from real data and random data. **a.**The plot shows the relationship of total number of SGAs being designated as SGA-FIs with respect to the threshold of calling an SGA-FI in random and real data. The x-axis shows the different thresholds, i.e., the number of DEGs predicted to be regulated by an SGA-FI, and the y-axis shows the number of significant SGA-FIs across all tumors, **b.** The plot shows the relationship of average number of SGAs being designated as SGA-FIs in a tumor with respect to the threshold of calling an SGA-FI in random and real data. The x-axis shows the different thresholds, i.e., the number of DEGs predicted to be regulated by an SGA-FI, and the y-axis shows the average number of significant SGA-FIs in a single tumor.

**Supplementary Figure S2.**
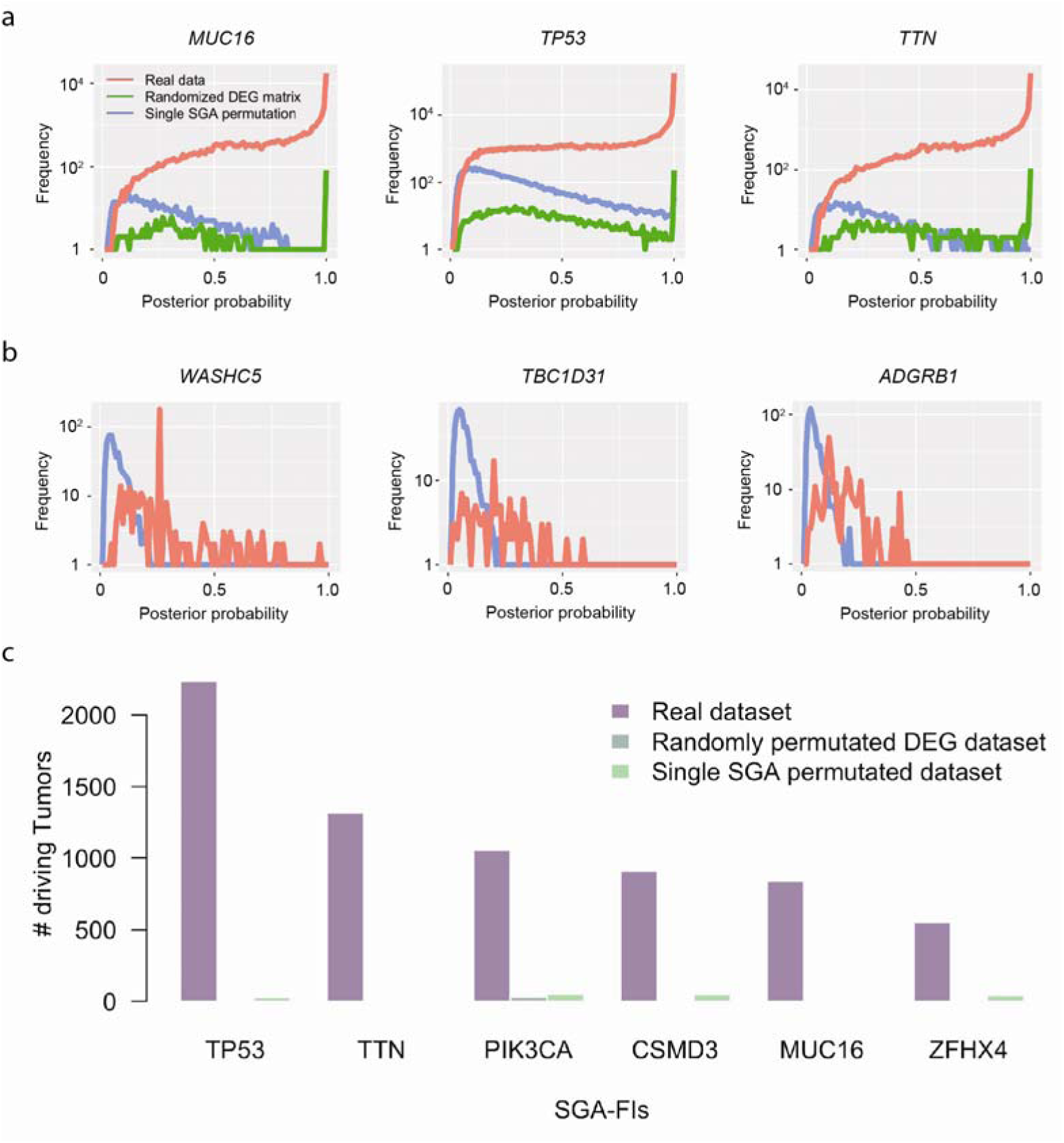
Comparison of causal analysis results from real data and random data. **a.**Comparison of distributions of the posterior probabilities of the highest candidate causal edges point from 3 most frequent SGAs to DEGs. **b.** Examples of 3 genes with high SGA frequency but without any high posterior probability causal edges emitting from them. **c.** Comparison of number of tumors called as SGA-FIs from the real dataset, randomly permutated DEG dataset and single SGA permutated dataset for the 6 most frequency SGAs.

**Supplementary Figure S3.**
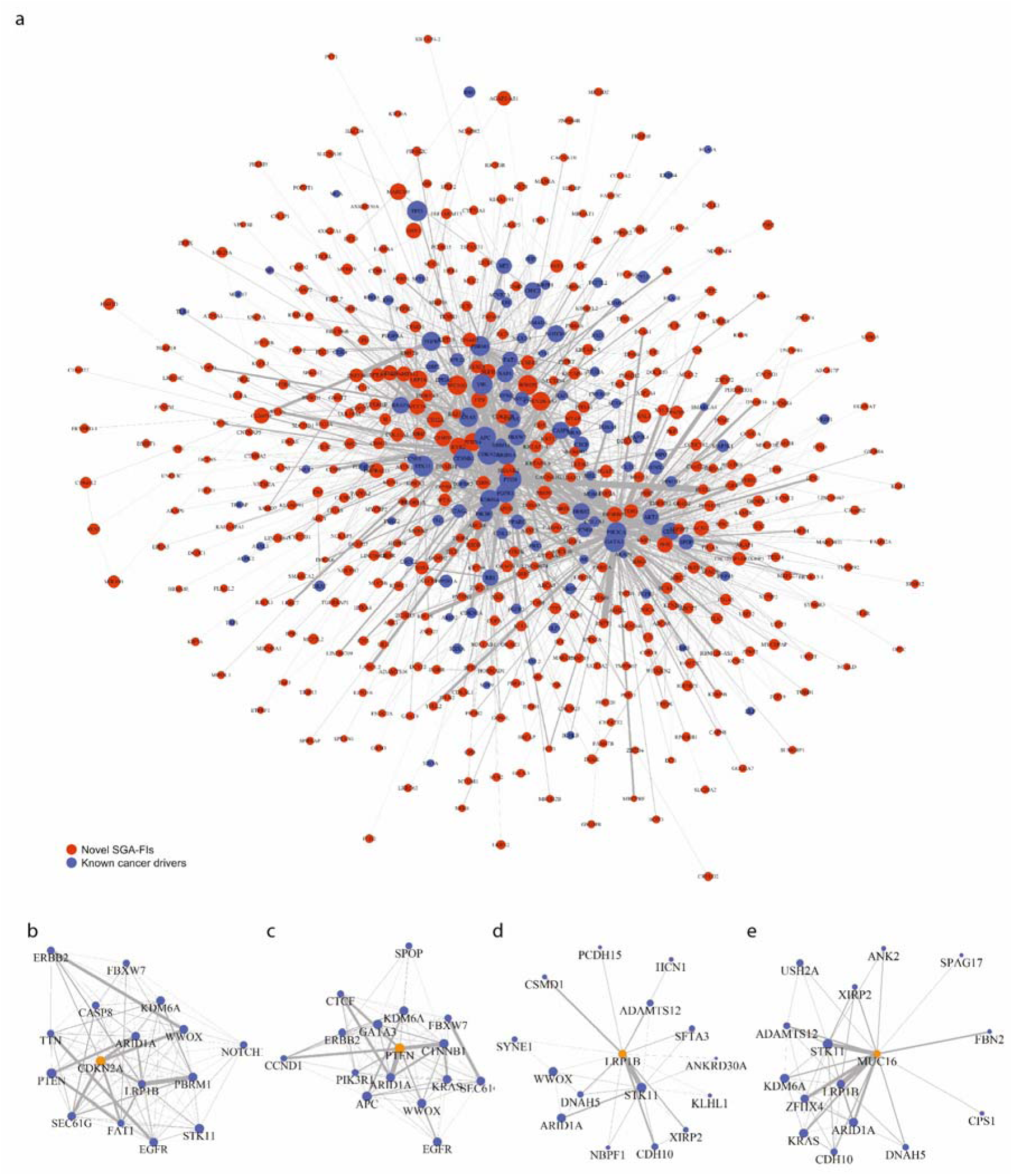
Networks of SGA-FIs share significant overlapping DEGs. **a.**SGA-FIs interacting network containing 536 SGA-FIs and 2669 edges. Blue nodes represent known cancer drivers and red nodes represent novel SGA-FIs. Node size indicates the number of its affected DEGs and edge width indicates the number of overlapped DEGs between two nodes, **b-e.** Top 15 SGA-FIs that share the most significant overlapping target DEGs with *CDKN2A, PTEN, LRP1B,* and *MUC16.* An edge between a pair of SGA-FI indicates that they share significantly overlapping target DEG sets, and the thickness of the line is proportional to negative log of the *p-*values of overlapping target DEG sets.

## Footnotes

1 The *call rate* for an SGA *A*_*h*_ is the ratio of number of tumors in which *A*_*h*_ is designated an SGA-FI over the number of tumors in which *A*_*h*_ occurs.

